# Immunization with a novel RNA replicon vaccine confers long-lasting protection against H5N1 avian influenza virus in 24 bird species

**DOI:** 10.1101/2025.01.15.633174

**Authors:** Marion Stettler, Stefan Hoby, Christian Wenker, Fabia Wyss, Elisabeth Heiderich, Lisa Butticaz, Nicolas Ruggli, Karin Darpel, Gert Zimmer

## Abstract

Highly pathogenic avian influenza viruses (HPAIV) of subtype H5N1 (clade 2.3.4.4b) have spread worldwide and caused the death of hundreds of millions of wild birds and domestic poultry. Moreover, spill over of H5N1 HPAIV from infected birds to more than 50 different mammalian species including humans has been recorded. While, licensed vaccines for protection of avian or mammalian species are not yet available, a few candidate vaccines are being trialled. Here, we report on the experimental vaccination of chickens and captive wild birds using a propagation-defective vesicular stomatitis virus (VSV), in which the essential envelope glycoprotein (G) protein gene was replaced by a modified hemagglutinin gene derived from a clade 2.3.4.4b H5N1 isolated in 2022 in the animal park of Bern, Switzerland. VSVΔG(H5_mb_) was produced on helper cells providing the VSV G protein *in trans*. Specific pathogen-free (SPF) chickens that were immunized twice via the intramuscular route with adjuvant-free VSVΔG(H5_mb_) replicon particles induced high levels of virus-neutralizing serum antibodies and were fully protected against lethal infection by H5N1 HPAIV (clade 2.3.4.4b). Notably, immunized animals did not shed challenge virus from the respiratory or gastrointestinal tract, suggesting that herd immunity can be achieved. The same vaccine was used to immunize a total of 317 captive wild birds at Bern Animal Park and Zoo Basel, representing 24 different species. No vaccine-associated side effects were observed. Birds without previous contact to H5Nx viruses produced high to very high H5-specific neutralizing antibody titers following the second immunization, while birds showing H5-specific antibodies prior to vaccination, already developed high neutralising antibody titers after a single immunization. One year after vaccination, most animals still showed significant neutralizing antibody titers, indicating that VSVΔG(H5_mb_) is able to induce a long-lasting protective immune response. Our results indicate that VSVΔG(H5_mb_) is an extraordinary safe and highly efficacious vaccine to stop H5N1 replication in various avian species.

## Introduction

Influenza A viruses belong to the family *Orthomyxoviridae* and are characterized by a single-stranded, negative-sense RNA genome consisting of 8 distinct segments ^1^. Influenza A viruses are further classified based on the two major antigens of the viral envelope, hemagglutinin (HA) and neuraminidase (NA). Nineteen, phylogenetically and serologically distinct HA subtypes and nine NA subtypes are known ^1–4^, and most of these subtypes have been detected in wild birds. In particular waterfowl is believed to represent the natural reservoir for avian influenza A viruses (AIV) ^5^. AIV normally do not cause any disease in these hosts and are therefore referred to as low-pathogenic avian influenza viruses (LPAIV). LPAIV predominantly replicate in the gastrointestinal tract of waterfowl and are shed along with the feces into the environment at large amounts. Sporadically, spill-over of LPAIV from wild birds to domestic poultry or mammalian species occurs.

The HA protein of AIV is synthesized as a precursor protein that needs to be proteolytically cleaved in order to perform low pH-triggered membrane fusion. Most avian AIV possess a monobasic cleavage site that is recognized by cellular trypsin-like proteases, which are expressed only in certain tissues. Viruses encoding HAs with a monobasic cleavage site cause only localized infections and usually display a low-pathogenic phenotype in most avian species ^6^. However, the HA may mutate and acquire additional basic amino acid residues at the cleavage site which allow recognition by ubiquitously expressed cellular prohormone convertases such as furin ^6,7^. AIV with such a polybasic cleavage site usually cause a systemic disease in poultry which is associated with up to 100 % mortality and known as “classical fowl plague”, a notifiable epizootic disease ^8^. Only highly pathogenic avian influenza viruses (HPAIV) of subtypes H5 and H7 have been detected in nature so far ^7^.

HPAIV of subtype H5N1was isolated for the first time from infected chickens in Scotland in 1959, and later was found to be responsible for an outbreak in turkeys in England in 1991 ^9^. In 1996, an outbreak by H5N1 HPAIV occurred in goose farms in Guangdong Province, China, and since then has spread globally. The Guangdong lineage diverged into distinct clades and subclades through mutation, reassortment and natural selection, and also expanded into multiple reservoir hosts ^10^. H5N1 HPAIV became endemic in several Asian countries and in Egypt. H5N1 HPAIV repeatedly spilled over to humans. From January 2003 to November 2024, 939 human cases of infection by H5N1 HPAIV of distinct clades have been reported by 24 countries, of which 464 cases were fatal ^11^. In 2020, a novel H5N1 HPAIV (clade 2.3.4.4b) emerged in the Middle East and became the most predominant variant in Asia, Europe, and Africa by 2021 ^12^. At the end of 2021, H5N1 of clade 2.3.4.4b was further disseminated by migratory birds from Europe to North America ^13^. Shortly thereafter, it reached Central and South America and finally found its way to Antarctica ^14^. At the beginning of 2024, viruses belonging to clade 2.3.4.4b have invaded all continents with the exception of Oceania. In the course of this panzootic, numerous wild birds succumbed to infection by H5N1 HPAIV with the consequence that populations of endangered avian species were reduced in some areas to levels that posed a threat to biodiversity and species conservation ^14–19^. In addition, H5N1 class 2.3.4.4b repeatedly caused mass deaths on poultry farms, with millions of birds succumbing to infection or had to be culled to contain the outbreaks.

H5Nx viruses of clade 2.3.4.4b have demonstrated a remarkable potential to infect a wide range of terrestrial and marine mammalian species ^20^, including foxes ^21,22^, minks ^23,24^, polar bears ^25^, cats ^26–28^, marine mammals ^14,29,30^, rodents ^31^ and dairy cows ^26,32,33^. Sequence analysis of viruses isolated from infected mammals revealed that H5N1 acquired mutations that were indicative of adaptation to mammals ^34^. H5N1 of clade 2.3.4.4b were also repeatedly transmitted to humans ^33–35^. In particular farm workers exposed to infected dairy cows and persons in charge of culling infected poultry were affected ^36,37^, but mostly presented only mild symptoms such as conjunctivitis ^38^. There is so far no evidence of human-to-human H5N1 transmission, however, the more the virus is spreading and adapts to mammalian hosts ^35^, the higher the pandemic potential of these viruses.

Zoological institutions represent a unique environment where several avian and mammalian species live in close proximity. Several bird species are kept in open pond systems that attracting wild birds looking for suitable resting and roosting sites and food sources. As a result, several zoos have reported outbreaks in the course of the panzootic by H5N1 HPAIV (clade 2.3.4.4b). Zoo birds are usually not vaccinated against H5N1 HPAIV and therefore are not protected. Due to the non-vaccination policy in the past, licensed vaccines are currently not available. Animal health requirements (e.g. housing indoor) often conflict with animal welfare requirements and lead to husbandry-related diseases if measures are imposed over longer periods of time. Experimental vaccination of zoo birds against H5N1 HPAIV using inactivated H5N2, H5N3 or H5N9 AIV revealed that captive wild bird species responded differentially to vaccination ^39–41^. Moreover, differentiation of infected from vaccinated animals (DIVA) remained a challenge, although the vaccine strains harbored an NA subtype other than N1. Thus, there is a need for safe and efficacious DIVA vaccines to protect avian as well as mammalian species from infection by H5N1. Such vaccines not only will protect animals from H5N1 infection, but will also reduce the risk of zoonotic H5N1 HPAIV transmission to humans.

Previous research has demonstrated that HA-recombinant single-cycle virus replicon particles (VRPs) based on vesicular stomatitis virus (VSV) can mediate effective protection of chickens from lethal infection by HPAIV of subtypes H7N1 ^42^ and H5N1 ^43^. These VRP vaccines elicited antibodies with neutralizing activity against other H5N1 clades, eliminated the shedding of challenge virus, and complied with the DIVA principle ^43^. In the present study, a VSV vector vaccine based on the HA antigen of H5N1 HPAIV (clade 2.3.4.4b) was used to immunize various bird species in two Swiss zoos. We evaluated and compared the level and duration of the humoral immune responses across different bird species over a 400-days period.

## Results

### Generation of a propagation-defective RNA replicon particle vaccine

In February 2022, an outbreak of H5N1 HPAIV occurred in Bern Animal Park, Switzerland, with two infected wild grey herons (*Ardea cinerea*) and one captive Dalmatian pelican (*Pelecanus crispus*) affected. Infectious virus was successfully isolated from biopsy material taken from the infected pelican. The viral RNA segments 4 (encoding HA) and 6 (encoding NA) of this virus isolate, which was designated A/Pelican/Bern/1/2022 (H5N1), were completely deciphered and submitted to the GISAID data bank (accession numbers EPI3526757 and EPI3526758, respectively). Comparison of the HA primary sequence with other published H5N1 sequences indicated that the virus belongs to the phylogenetic clade 2.3.4.4b.

In previous work, we showed that SPF chickens that were immunized via the intramuscular route with VSVΔG(HA) replicon particles encoding the HA gene of A/chicken/Yamaguchi/7/2004 (H5N1), clade 2.5, were fully protected from lethal infection with HPAIV A/whooper swan/Mongolia/3/2005 (H5N1) belonging to clade 2.2 ^43^. We compared the primary amino acid sequence of the HA_1_ subunit of A/Pelican/Bern/1/2022 (H5N1) with those of previous H5 clades (**Supplementary Fig. 1**). We found that the HA_1_ subunit of this clade 2.3.4.4b virus differed in several amino acid positions from the HA_1_ subunits of older H5 clades. Of note, most of these substitutions clustered to the HA globular head region which harbors the receptor binding domain.

In order to generate a vaccine perfectly matching the HA of currently circulating clade 2.3.4.4b H5N1, we generated VSVΔG(H5) replicon particles encoding either the authentic HA gene of A/Pelican/Bern/1/2022 (H5N1) with the typical polybasic (pb) HA cleavage site or encoding a modified HA in which the polybasic cleavage site was changed into a monobasic (mb) sequence motif typically found in low-pathogenic H5 viruses (**Fig. 1a**). The replicon particles were propagated on baby hamster kidney cells conditionally expressing the VSV G protein ^44^, resulting in 1.7 x 10^8^/mL of infectious particles at 20 hours post infection (p.i.) (**Fig. 1b**). The virus replicon particles (VRP) were pelleted from the infected cell culture supernatant by ultracentrifugation and resuspended in a tenfold smaller volume of PBS. Titration of these vaccine stocks revealed titers of 8.5 x 10^8^ f.f.u./mL (**Fig. 1b**), indicating that this purification step resulted in a 5-fold concentration of infectious replicon particles.

**Fig. 1.**
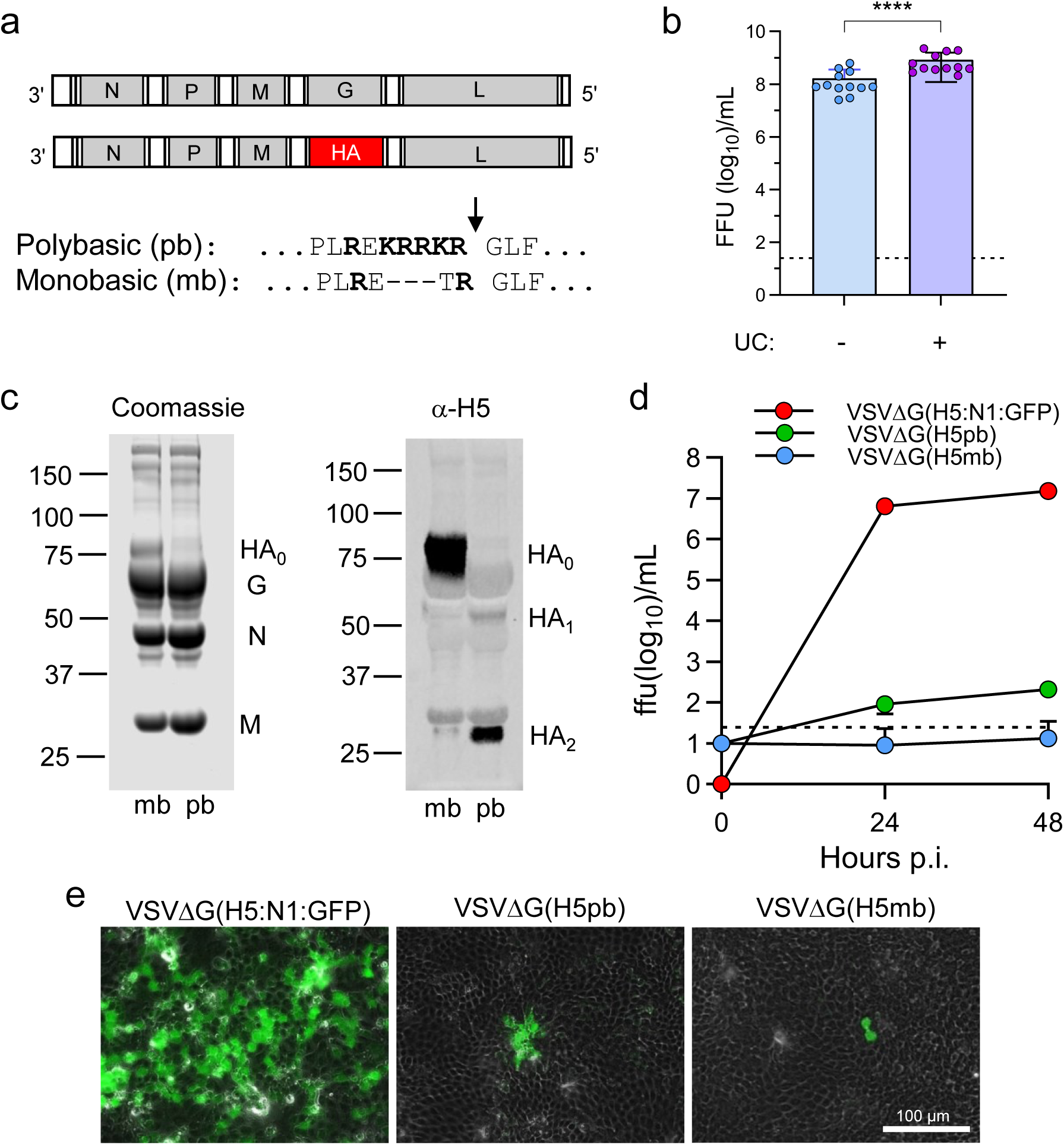
Generation and characterization of VSVΔG(H5) replicon particles. (**a**) Genome maps of authentic VSV and G-deficient VSVΔG(H5) vector. Authentic VSV contains five transcription units encoding the N, P, M, G, and L gene, respectively. In VSVΔG(H5), the G gene was replaced with the HA gene of A/Pelican/Bern/1/2022 (H5N1) encoding either the original polybasic (pb) or a monobasic (mb) proteolytic cleavage site. (**b**) Infectious virus yield on *trans*-complementing helper cells. BHK-G43 cells were treated with mifepristone (MFP) to trigger expression of the VSV G protein, or were left untreated prior to infection with VSVΔG(H5_mb_) using 0.1 f.f.u./cell. The cell culture supernatant (200 mL) was harvested at 20 hours p.i., virus particles concentrated by ultracentrifugation (UC) at 105’000 x g, and resuspended in 20 mL of PBS. Infectious virus titers before and after ultracentrifugation of the cell culture supernatant were determined on BHK-21 cells. Mean values and standard deviations of 12 vaccine batches are shown. The one-way ANOVA test was used to analyze whether ultracentrifugation has led to significantly higher infectious titers (****p < 0.0001). (**c**) Western blot analysis of VSVΔG(H5_pb_) and VSVΔG(H5_mb_) particles propagated on MFP-treated BHK-G43 helper cells and concentrated by ultracentrifugation. Pelleted virus particles were dissolved in SDS sample buffer containing 0.1 M dithiothreitol and viral proteins separated by SDS-PAGE. The protein composition of the virions was visualized by staining with colloidal Coomassie (left panel), or after transfer to nitrocellulose membrane by immunostaining using a chicken polyclonal anti-H5 serum. (**d**) Multi-cycle virus replication on MDCK cells. Cells were infected with 0.0001 infectious virus particles per cell, and cell culture supernatant sampled at 1, 24, and 48 hours post infection. Infectious titers were determined on BHK-21 cells. Mean values and standard deviations of 3 infection experiments are shown. The lower limit of detection (25 f.f.u./mL) is represented by a dashed line. (**e**) Immunofluorescence analysis of MDCK cells 20 hours p.i. with the indicated viruses (0.02 infectious units per cell). Cells infected with VSVΔG(H5_P_:N1_P_:GFP) were detected taking advantage of the GFP reporter protein. Cells infected with either VSVΔG(H5_mb_) or VSVΔG(H5_pb_) were detected by indirect immunofluorescence using an NP-specific monoclonal antibody.

Analysis of the purified replicon particles by SDS gel electrophoresis under reducing conditions and subsequent staining with colloidal Coomassie revealed the major constituents of VSV, i.e. nucleoprotein (N), envelope glycoprotein (G), and matrix (M) protein (**Fig. 1c, left panel**). An additional band corresponding to the relative molecular mass of 80 kDa was detected in VSVΔG(H5_mb_) particles. This band was recognized in Western blot analysis by a H5N1-specific polyclonal chicken serum (**Fig. 1c, right panel**), and likely represents the non-cleaved HA_0_ precursor. In accordance with the processing of authentic HA in the *trans*-Golgi network by the prohormone convertase furin, the HA_0_ precursor was not present in purified VSVΔG(H5_pb_) particles which rather contained the cleavage products HA_1_ and HA_2_. These findings confirmed previous results which showed that recombinant HA is incorporated into the VSV envelope, albeit at much lower levels compared to the VSV G protein ^45^.

To assess the replication competence of VSVΔG(H5), multi-cycle virus replication kinetics were performed on Madin-Darby canine kidney (MDCK) cells using a multiplicity of infection (m.o.i.) of 0.0001 infectious virus particles per cell. Titration of infectious virus particles released into the cell culture supernatant revealed that VSVΔG(H5_mb_) harboring a monobasic HA cleavage site did not produce any infectious particles (**Fig. 1d**) and remained restricted to single infected cells (**Fig. 1e**). VSVΔG(H5_pb_) showed a limited cell-to cell spread in MDCK monolayers (**Fig. 1d**), with some infectious particles released into the cell culture medium (**Fig. 1e**). In contrast, VSVΔG(H5_pb_:N1:GFP) encoding the authentic HA and NA antigens of A/Pelican/Bern/1/2022 (H5N1) replicated in MDCK cells in an autonomous manner, reaching about 10^7^ f.f.u./mL at 48 hours p.i. (**Fig. 1d**). In line with a previous report, this finding demonstrates that NA is essentially required for the release of infectious viral particles ^46^. Because of the even more restricted release of VSVΔG(H5_mb_) compared to VSVΔG(H5_pb_) (**Fig. 1d, e**), the VSVΔG(H5_mb_) vaccine was selected for further evaluation in animals.

### Chickens vaccinated with VSVΔG(H5_mb_) were fully protected from lethal challenge with highly pathogenic H5N1 (clade 2.3.4.4b)

The VSVΔG(H5_mb_) vaccine was evaluated with 5-weeks old specific pathogen-free (SPF) chickens (White Leghorn). The animals were divided in 4 groups each containing 8 chickens (mixed sex). Animals of group A were immunized via the intramuscular (i.m.) route with VSV*ΔG, a VRP encoding the green fluorescent protein (GFP) but not any influenza antigen (**Fig. 2a**). Chickens of animal groups B and C were immunized with VSVΔG(H5_mb_) via the intramuscular route, while chickens of animal group D were immunized with the same vaccine via the ocular route. At day 28, groups A and B were boosted (i.m.) with VSV*ΔG and VSVΔG(H5_mb_), respectively, while chickens of groups C and D were booster vaccinated with VSVΔG(H5_mb_) via eye drop. At day 56, chickens of all groups were infected via the nasal route with 10^6^ TCID_50_ of A/Pelican/Bern/1/2022 (H5N1).

**Fig. 2.**
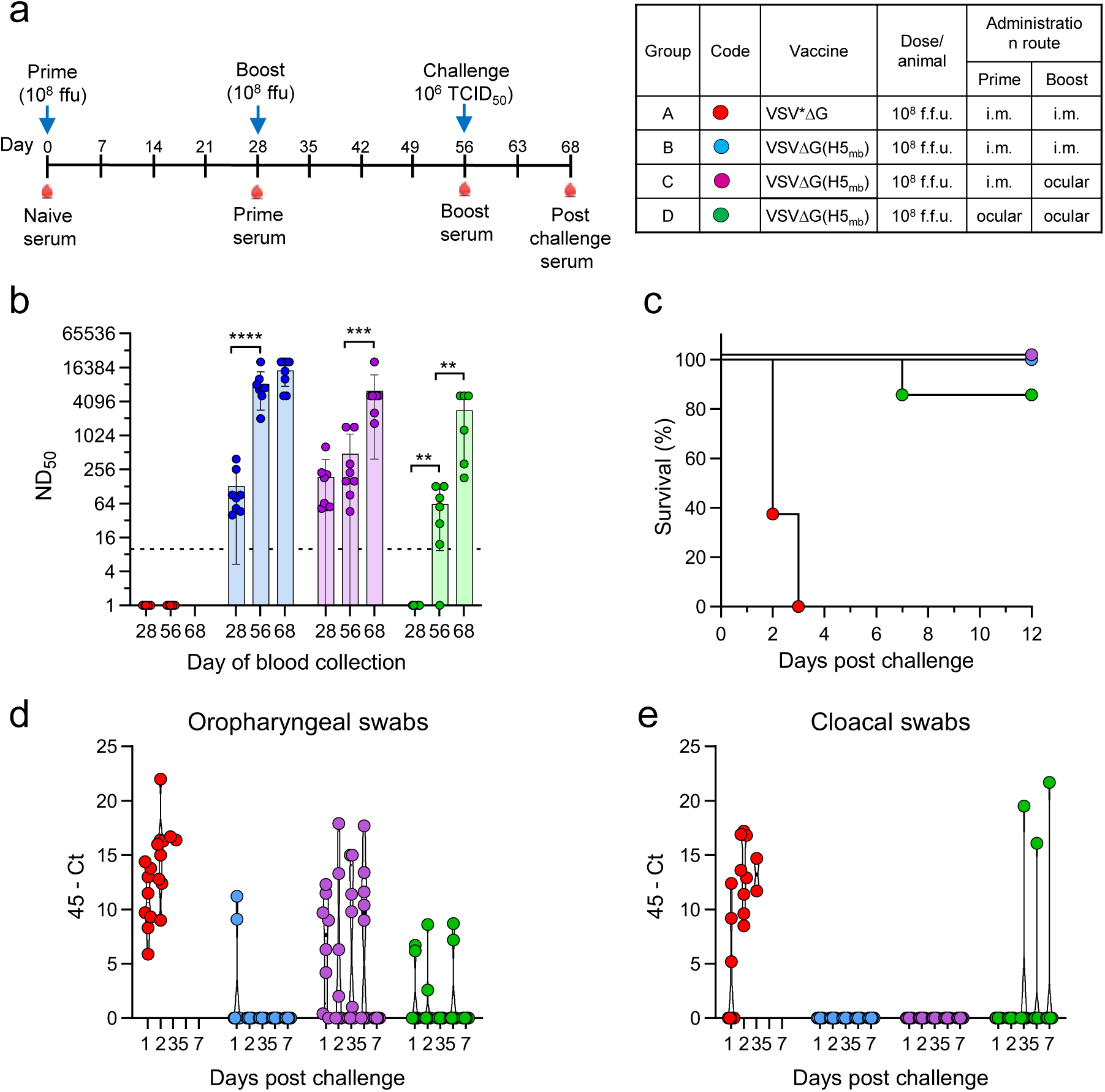
Evaluation of the VSVΔG(H5_mb_) vaccine in specific pathogen-free chickens. (**a**) Schematic representation of the experimental design. Red drops below the timeline indicate time points of blood sampling. The four vaccine groups (8 chickens per group) with the corresponding color code, vaccine name, dose, and administration route are depicted on the right-hand side. (**b**) Determination of the virus neutralization dose 50% (ND_50_) in serum of immunized animals. Mean values and standard deviations (SD) are shown. The two-way ANOVA test was used to assess significant ND_50_ values between prime, boost and post-challenge sera (**p < 0.01, ***p < 0.001, ****p < 0.0001). (**c**) Survival rate of SPF chickens following nasal infection with 10^6^ TCID_50_ of A/Pelican/Bern/1/2022 (H5N1). (**d, e**) Determination of virus load by RT-qPCR in oropharyngeal (**d**) and cloacal (**e**) swab samples collected at the indicated day p.i. with H5N1 HPAIV.

Immune sera were analysed for virus-neutralizing activity using VSVΔG(H5_P_:N1_P_:GFP), a propagation-competent chimeric virus encoding the HA and NA of A/Pelican/Bern/1/2022 (H5N1) and a GFP reporter. This surrogate virus allowed us not only to perform virus-neutralisation test at biosafety level 2, the GFP reporter also facilitated the readout of the assay as cells non-protected from infection were easily and rapidly detected. We found that the neutralisation dose 50% (ND_50_) that was determined with this surrogate virus almost perfectly matched the ND_50_ values that were obtained with the authentic HPAIV A/Pelican/Bern/1/2022 (H5N1) (**Supplementary Fig. 2**).

Chickens of group A which had been vaccinated with the control vaccine VSV*ΔG did not develop antibodies with neutralizing activity against VSVΔG(H5_P_:N1_P_:GFP) (**Fig. 2b)**. The animals of this control group were not protected from H5N1 HPAIV challenge infection and died at day 2 or day 3 p.i. (**Fig. 2c**). The H5N1 HPAIV challenge virus was shed from the oropharyngeal and cloacal site of group A animals already starting one day after infection. Virus shedding continued until the animals succumbed to infection or had to be euthanized due to severe symptoms (**Fig. 2d**). Analysis of the immune sera that were prepared from groups B and C four weeks after the first immunization with VSVΔG(H5_mb_) revealed significant levels of neutralizing antibodies (nAb) (mean ND_50_ values of 130 and 185, respectively). After the second immunization with VSVΔG(H5_mb_), nAb titers increased 64-fold in group B (mean ND_50_ value of 8347), whereas no significant increase of nAb was observed when group C chickens were immunized at day 28 via the ocular route. All animals in these two vaccine groups survived nasal infection by HPAIV A/Pelican/Bern/1/2022 (H5N1) (**Fig. 2c**), without showing any symptoms of disease (data not shown).

RT-qPCR analysis of swab samples collected from each animal at days 1, 2, 3, 5 and 7 p.i. demonstrated that group A animals shed high levels of H5N1 challenge virus from both the oropharyngeal (**Fig. 2d**) and cloacal (**Fig. 2e**) site. Two animals in group B showed oropharyngeal H5N1 shedding at day 1 p.i. only, while shedding from the cloacal site was not detected (**Fig. 2d**). Interestingly, cloacal but not oropharyngeal shedding of H5N1 HPAIV was inhibited in group C. Immune sera prepared at day 12 p.i. showed no increase of nAb titers in group B (**Fig. 2b**). However, in group C titers increased significantly, indicating that the challenge virus was still able to replicate in the vaccinated animals thereby boosting the immune response.

Group D chickens did not develop significant nAb levels after the first ocular immunization, but developed significant nAb titers when the vaccine was administered the second time via the same route (mean ND_50_ value of 62). With the exception of one animal, which did not develop any detectable nAbs, all animals of this group survived challenge infection with H5N1 HPAIV (**Fig. 2c**), however, some animals still shed H5N1 HPAIV from the oropharyngeal and/or cloacal site (**Fig. 2d, e**). Similar to group C, challenge infection largely boosted the immune response in the vaccinated animals. Together, these findings indicate that the prime/boost vaccination regimen using VSVΔG(H5_mb_) VRP induced a robust and protective immune response that largely blocked H5N1 HPAIV shedding.

### Some avian species had pre-existing immunity to H5Nx

In order to assess the immune status of zoo birds prior to vaccination, sera were prepared from all animal groups and tested by ELISA for the presence of antibodies against influenza nucleoprotein (NP), a highly conserved antigen shared by different influenza subtypes. We found that the majority of the 24 avian species tested were negative for NP-specific antibodies (**Tab. 1)**. However, all the sera that had been collected from Greater flamingos (*Phoenicopterus roseus*) in the animal park of Bern and the zoo of Basel were tested positive for NP-specific antibodies, indicating that they had been infected in the past with AIV of unknown subtype and pathotype. Likewise, some or all birds of the Eastern White pelicans (*Pelecanus onocrotalus*), common Eider ducks (*Sommateria mollissima*), Barnacle geese (*Branta leucopsis*), Nene geese (*Branta sandvicensis*), Bar-headed geese (*Anser indicus*), Black swans (*Cygnus atratus*), and Coscoroba swans (*Coscoroba coscoroba*) were found positive for NP-specific antibodies. Interestingly, all these avian species were kept in pond systems frequently visited by wild ducks and grey herons among others. Sera that had been tested positive for NP-specific antibodies were subsequently tested for the presence of H5-specific antibodies (**Tab. 1**). It was found that all NP-positive Greater flamingos (*Phoenicopterus roseus*) in Bern Animal Park and a large fraction of the other NP-positive birds also had H5-specific antibodies, indicating that these birds must have been infected with subtype H5 AIV. However, it was not possible to determine by ELISA whether these have been low-pathogenic or highly pathogenic H5Nx viruses.

**Table 1.** Serological (ELISA) analysis of avian species enrolled in the vaccination program.

| Species | Common name | Site | Animals | NP<sup>+</sup> (d1) | H5<sup>+</sup> (d1) | NP<sup>+</sup> (d365) |
| --- | --- | --- | --- | --- | --- | --- |
| *Gallus gallus dom.* | Orpington chicken | Basel | 3 | 0/3 | ND | 0/2 |
|  | Silky chicken | Basel | 32 | 0/32 | ND | 0/27 |
|  | Appenzeller Spitzhaube chicken | Bern | 5 | 0/4 | ND | 0/5 |
| *Anser anser dom.* | Diepholzer goose | Bern | 2 | 0/2 | ND | 0/2 |
| *Pyrhacorax pyrrhacorax* | Chough | Bern | 2 | 0/2 | ND | 0/2 |
| *Ciconia nigra* | Black stork | Bern | 2 | 0/2 | ND | 0/2 |
| *Bubo bubo* | Eurasian eagle owl | Bern | 2 | 0/2 | ND | 0/2 |
| *Aegolius funereus* | Boreal owl | Bern | 2 | 0/2 | ND | 0/1 |
| *Athene noctua* | Little owl | Bern | 2 | 0/2 | ND | 0/1 |
|  |  | Basel | 2 | 0/2 | ND | 0/1 |
| *Nyctea scandiaca* | Snowy owl | Bern | 2 | 0/2 | ND | 0/2 |
| *Tetrao urogallus* | Western capercaillie | Bern | 7 | 0/7 | ND | 0/3 |
| *Alectoris graeca* | Rock partridge | Bern | 8 | 0/8 | ND | 0/5 |
| *Phoenicopterus roseus* | Greater flamingo | Bern | 66 | 33/33 | 33/33 | 33/33 |
|  |  | Basel | 101 | 54/54 | 45/45 | 52/52 |
| *Pelecanus crispus* | Dalmatian pelican | Bern | 10 | 1/10 | 1/1 | 1/10 |
|  |  | Basel | 8 | 0/8 | ND | 0/7 |
| *Pelecanus onocrotalus* | Eastern white pelican | Basel | 19 | 10/19 | 7/10 | 11/19 |
| *Spheniscus demersus* | African penguin | Basel | 45 | 0/45 | ND | 1/45 |
| *Aythya nyroca* | Ferruginous duck | Bern | 3 | 0/3 | ND | 0/3 |
| *Bucephala clangula* | Common goldeneye | Bern | 1 | 0/1 | MD | ND |
| *Somateria mollissima* | Common Eider duck | Bern | 4 | 3/4 | 1/3 | 3/4 |
| *Branta ruficollis* | Red-breasted goose | Basel | 4 | 3/4 | ND | 2/2 |
| *Branta leucopsis* | Barnacle goose | Basel | 2 | 1/2 | 0/1 | 0/2 |
| *Branta sandvicensis* | Nene goose | Basel | 4 | 4/4 | 0/4 | 1/4 |
| *Anser indicus* | Bar-headed goose | Basel | 2 | 1/2 | 0/1 | 0/2 |
| *Anser anser* | Greylag goose | Bern | 4 | 0/4 | ND | 3/3 |
| *Cygnus atratus* | Black swan | Basel | 2 | 2/2 | 2/2 | 2/2 |
| *Coscoroba coscoroba* | Coscoroba swan | Basel | 2 | 1/2 | 1/1 | 2/2 |
| **Σ = 24** |  |  | **Σ = 348** |  |  |  |

ND: Not determined.

### Vaccinated zoo birds produce high levels of long-lasting H5N1-neutralizing antibodies

A total of 317 birds, belonging to 24 species were immunized via the intramuscular route with adjuvant-free VSVΔG(H5_mb_) suspension. The animals received a second dose of the same vaccine 35 days after the primary immunisation and a third injection at day 365 in order to boost the immune response after one year (**Fig. 3a**). No obvious adverse effects due to the vaccination were observed, indicating that the vaccine was well tolerated. Blood was sampled at days 1, 35, 70, 365, and 400, sera prepared, and analysed for virus-neutralizing activity using the VSVΔG(H5_P_:N1_P_:GFP) reporter virus. Interestingly, the Greater flamingos and Black swans displayed high levels of virus-neutralizing activity in pre-immune serum, while in most other species nAb were not detected or were found at low levels (**Tab. 2**). Almost all animals responded to the first immunisation with VSVΔG(H5_mb_) by formation of nAb at levels that are regarded as protective (ND_50_ ≥ 10) (**Fig. 3**, **Tab. 2**). However, there were notable differences between species in the level of the immune response. In particular those species that showed virus-neutralizing activity prior to vaccination, e.g. Greater flamingos, Red-breasted geese and Black swans, responded to the first immunization by producing high nAb titers, suggesting that the primary immunisation with VSVΔG(H5_mb_) boosted pre-existing immunity to H5 in these animals. When these animals were immunized the second time, ND_50_ titers increased only moderately, while in avian species with no pre-existing immunity to H5N1 nAb levels increased significantly in response to the second immunisation (**Fig. 3**, **Tab. 2**). Silky chickens, for example, had no nAbs prior to vaccination, developed a mean ND_50_ titer of 68 after the first immunisation, and reached a mean ND_50_ titer of 1239 after the second immunisation (**Tab. 2**). Analysis of day 365 and day 400 sera of immunized Silky chickens showed that nAb titers had dropped only moderately over one year, and increased slightly when the animals were boosted again at day 365 (**Fig. 3b**). Analysis of the day 365 sera by NP-ELISA showed that the chickens did not experience any AIV infection in this time (**Tab. 1**). Analysis of immune sera prepared from immunized African penguins (*Spheniscus demersus*) at days 70 and 365 showed that the mean ND_50_ value dropped from 745 to 309 (**Fig. 3c**, **Tab. 2**), a nAb titer considered as protective. The third immunisation again stimulated the immune response as indicated by mean ND_50_ titers of 1264 at day 400. Similar though not identical kinetics of virus-neutralizing antibody titers in response to repeated vaccination with VSVΔG(H5_mb_) were found for immunized Greater flamingos (**Fig. 3d, e**), the two Pelican species (**Fig. 3f**), as well as for most other avian species enrolled in the study (**Tab. 2**).

**Fig. 3.**
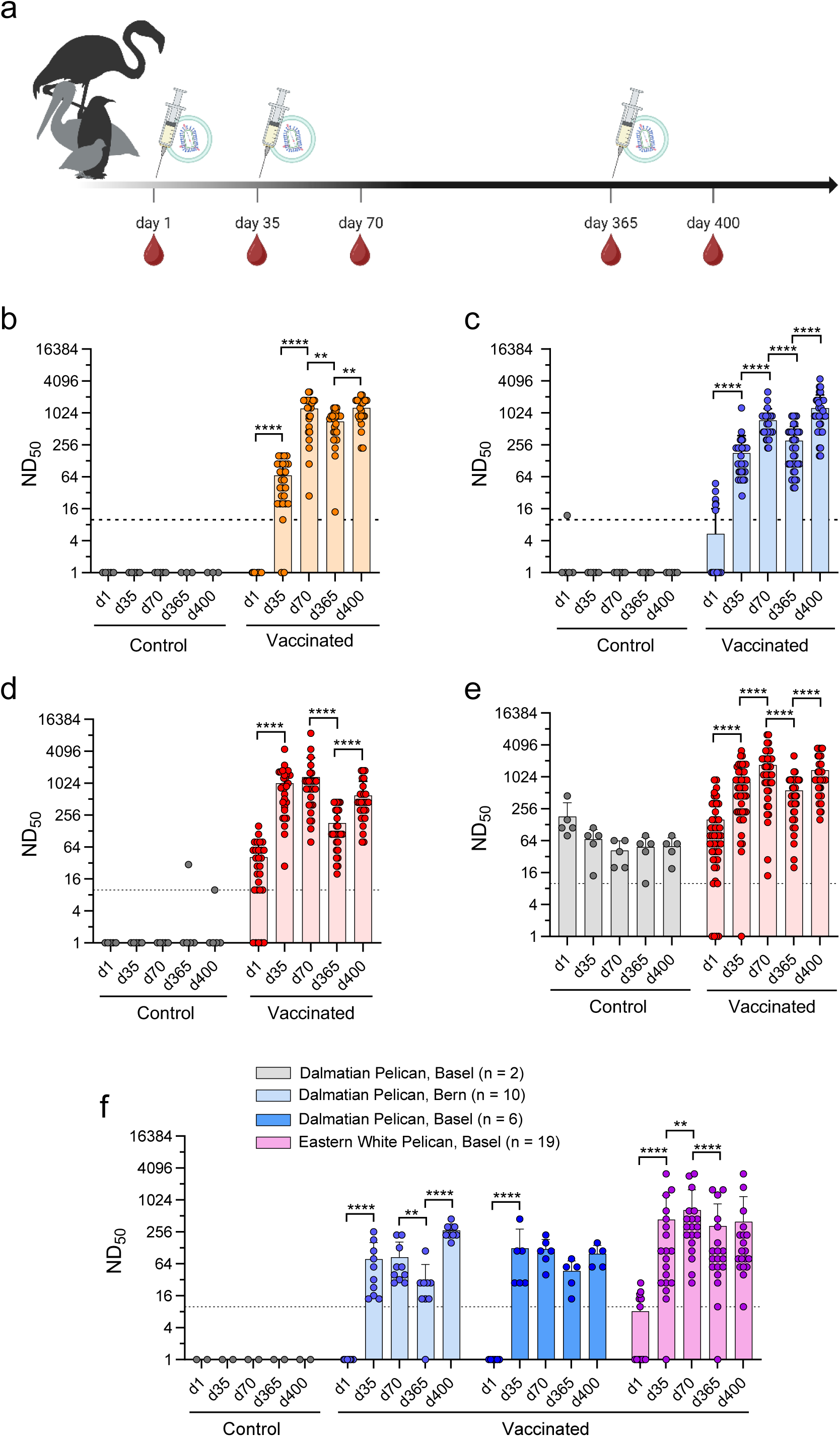
Evaluation of the VSVΔG(H5mb) vaccine in captive wild birds. (**a)** Study design. A total of 317 captive wild birds in Bern Animal Park and the zoo of Basel representing 24 different species were immunized intramuscularly with 2x10^8^ f.f.u./mL of VSVΔG(H5_mb_) using 250 μl (5x10^7^ f.f.u.) for small birds (< 1.5 kg of body weight) and 500 μl for large birds (≥ 1.5 kg of body weight). The animals were boosted at day 35 and day 365 using the same vaccine, dose and administration route. Blood samples were collected at days 1, 35, 365 and 400, and serum analyzed by virus-neutralization tests and ELISA. The figure was created using biorender.com. (**b-e)** Determination of H5-specific nAb titers in serum samples prepared from non-vaccinated controls and vaccinated birds at the indicated days. The VSVΔG(H5_P_:N1_P_:GFP) surrogate virus was used for the virus-neutralization tests. (**b**) Determination of nAb titers in immune sera of Silky chickens (*Gallus gallus domesticus*; n = 32). (**c**) Determination of ND_50_ titers in immune sera of African penguins (*Spheniscus demersus*; n = 45). (**d, e**) Detection of virus-neutralizing activity in sera of Greater Flamingos (*Phoenicopterus roseus*) living in Bern Animal Park (**d,** n = 33) and the zoo of Basel (**e,** n = 54). ND_50_ values of one animal were excluded due to their extraordinary high levels (d1: 44’668, d35: 64’000, d70: 44’668, d365: 64’000, d400: 89’125). (**f**) Determination of ND_50_ titers in sera of Dalmatian pelicans (*Pelecanus crispus*) living in Bern Animal Park (n = 10) and the zoo of Basel (n = 8), and in sera of Eastern White pelicans (*Pelecanus onocrotalus*) living in the zoo of Basel (n = 19). Mean values and standard deviations are shown. The lower detection limit (ND_50_ < 10) is indicated. Statistically significant ND_50_ values were computed using the two-way ANOVA (**b-e**) or the one-way ANOVA (**f**) with Tukey’s multiple comparison test (\*\**p* < 0.01, \*\*\*\**p* < 0.0001).

**Table 2.** Virus-neutralizing antibody titers in serum of zoo birds prior and post vaccination with VSVΔG(H5_mb_).

| Species | Common name | Location | Number of animals | ND<sub>50</sub> titer<br>in serum collected at day |  |  |  |  |
| --- | --- | --- | --- | --- | --- | --- | --- | --- |
|  |  |  |  | d1 | d35 | d70 | d365 | d400 |
| *Gallus gallus dom.* | Orpington chicken | Basel | 3 | <10 | 27<br>±13 | 330<br>±112 | ND, 40, 160 | ND, 160,<br>320 |
|  | Silky chicken | Basel | 27 | <10 | 68<br>±53 | 1239<br>±710 | 705<br>±375 | 1268<br>±637 |
|  | Appenzeller Spitzhaube chicken | Bern | 5 | <10 | 52<br>±32 | 797<br>±538 | 94<br>±81 | 281<br>±359 |
| *Anser anser dom.* | Diepholzer goose | Bern | 2 | 14, <10 | 65, 257 | 160, 320 | ND, 160 | ND, 1600 |
| *Pyrhacorax pyrrhacorax* | Chough | Bern | 2 | <10 | <10, 80 | 182, 1280 | 14, 80 | 4467, 2560 |
| *Ciconia nigra* | Black stork | Bern | 2 | <10 | 446, 363 | 128, 363 | 14, 20 | 224, 447 |
| *Bubo bubo* | Eurasian eagle owl | Bern | 2 | <10 | 80, 91 | 640, 1023 | 28, 14 | 447, 640 |
| *Aegolius funereus* | Boreal owl | Bern | 2 | <10 | ND, ND | 182, 320 | 20, ND | 640, ND |
| *Athene noctua* | Little owl | Bern | 2 | <10 | ND, 32 | ND, 1280 | ND, 447 | ND, 447 |
|  |  | Basel | 2 | <10, <10 | 14, 112 | 891, 3200 | 80, 160 | 320, 4467 |
| *Nyctea scandiaca* | Snowy owl | Bern | 2 | <10, <10 | 28, 20 | 724, 80 | 20, 14 | 447, 320 |
| *Tetrao urogallus* | Western capercaillie | Bern | 3 | <10 | 90<br>±70 | 611<br>±360 | 40<br>±35 | 2346 ±805 |
| *Alectoris graeca* | Rock partridge | Bern | 5 | <10 | 9<br>±9 | 471<br>±736 | 362<br>±309 | 1292 ±1804 |
| *Phoenicopterus roseus* | Greater flamingo | Bern | 28 | 41<br>±39 | 1023<br>±914 | 1334<br>±1760 | 181<br>±151 | 602<br>±511 |
|  |  | Basel | 48<sup>a</sup> | 160<br>±213 | 835<br>±752 | 1713<br>±1491 | 569<br>±469 | 1362<br>±1086 |
| *Pelecanus crispus* | Dalmatian pelican | Bern | 10 | <10 | 79<br>±83 | 86<br>±79 | 30<br>±32 | 274<br>±81 |
|  |  | Basel | 6 | <10 | 126<br>±163 | 121<br>±64 | 47<br>±26 | 99<br>±44 |
| *Pelecanus onocrotalus* | Eastern white pelican | Basel | 19 | 8<br>±10 | 439<br>±802 | 660<br>±924 | 328<br>±533 | 396<br>±788 |
| *Spheniscus demersus* | African penguin | Basel | 40 | 5<br>±11 | 179<br>±205 | 745<br>±482 | 309<br>±266 | 1264<br>±968 |
| *Aythya nyroca* | Ferruginous duck | Bern | 3 | <10 | 14 | 68<br>±17 | 25<br>±5 | 120<br>±91 |
| *Bucephala clangula* | Common goldeneye | Bern | 1 | <10 | 40 | 40 | 10 | 56 |
| *Sommateria mollissima* | Common Eider duck | Bern | 4 | <10 | 117<br>±137 | 1016<br>±505 | 95<br>±89 | 1401<br>±820 |
| *Branta ruficollis* | Red-breasted goose | Basel | 2 | 14, 14 | 2239, 1600 | 1600, 3200 | 447, 891 | 640, 1280 |
| *Branta leucopsis* | Barnacle goose | Basel | 2 | <10, 10 | 10, 20 | 20, 28 | 14, 28 | 56, ND |
| *Branta sandvicensis* | Nene goose | Basel | 4 | 15<br>±4 | 28<br>±22 | 543<br>±277 | 158<br>±119 | 1280, ND,<br>ND, 1778 |
| *Anser indicus* | Bar-headed goose | Basel | 2 | <10, 14 | 447, 20 | 447, 112 | 80, 28 | 447, 112 |
| *Anser anser* | Greylag goose | Bern | 4 | 3<br>±3 | 100<br>±24 | 255<br>±112 | 88<br>±42 | 469<br>±161 |
| *Cygnus atratus* | Black swan | Basel | 2 | 447, 56 | 891, 1280 | 1778, 1778 | 640, 891 | 640, 891 |
| *Coscoroba coscoroba* | Coscoroba swan | Basel | 2 | 20, <10 | 112, 28 | 112, 112 | 56, 40 | 447, 1778 |

ND: Not determined; <sup>a</sup> ND<sub>50</sub> values of one animal were excluded due to their extraordinary high levels (d1: 44'668, d35: 64'000, d70: 44'668, d365: 64'000, d400: 89'125).

### VSVΔG(H5_mb_) induces significant levels of VSV-neutralizing antibodies

As neutralizing antibodies directed to the VSV G protein might interfere with repeated administration of VRPs bearing the VSV G protein on their surface, we also tested immune sera of Dalmatian pelicans and Silky chickens for VSV-neutralizing activity (**Supplementary Fig. 3**). We did not detect significant neutralizing activity against VSV in sera of non-vaccinated Dalmatian pelicans and in pre-immune sera. In sera of day 35, some vaccinated Dalmatian pelicans and at day 70 all vaccinated pelicans showed significant levels of VSV-specific nAbs (**Supplementary Fig. 3a)**. At day 365, nAbs titers have declined significantly, with some being below the detection limit. After the annual booster immunization, VSV-specific nAbs increased again, indicating that the induction of VSV- and H5-specific nAbs in VSVΔG(H5_mb_)-immunized Dalmatian pelicans followed a similar kinetics (see **Fig. 2f**). This was also true for immunized Silky chickens, as both VSV-specific (**Supplementary Fig. 3b**) and H5-specific nAbs (**Fig. 3b**) showed similar titers on days 365 and 400 compared to day 70. These findings suggest that although VSVΔG(H5_mb_)-infected cells do not express VSV G protein, the amount of G protein on the surface of replicon particles is sufficient to elicit significant levels of VSV-neutralizing antibodies.

### VSVΔG(H5_mb_) particles elicit antibodies with broadly H5-specific neutralizing activity

We found that several amino acid residues in the globular head domain of H5 hemagglutinin (clade 2.3.4.4b) differed from those of older clades (see **Supplementary Fig. 1**). For this reason, the recombinant VRP vaccine used in this study was adapted to the antigenic features of the currently circulating panzootic H5N1. To assess whether the antibodies induced by the updated vaccine would inhibit previous H5 clades, we analyzed day 70 immune sera from Silky chickens and African penguins for neutralizing activity against VSVΔG(H5:N1:GFP) surrogate viruses encoding the HA antigen of either clade 1.1, clade 2.5, clade 2.3.2.1, and clade 2.3.4, and used a surrogate virus with the homotypic HA (clade 2.3.4.4b) as a reference (**Fig. 4**). Compared to the neutralizing activity against surrogate virus with homotypic HA, the mean ND_50_ values obtained with previous clades were significantly lower (**Fig. 4b**). However, all but one animal showed ND_50_ titers which can be considered as protective. Interestingly, the HA antigens of previous H5 clades, including clade 2.3.4, were only distantly related to the HA antigen of clade 2.3.4.4b. Likewise, day 70 immune sera from African penguins showed significantly lower mean ND_50_ titers against earlier clades when compared to homotypic clade 2.3.4.4b. Of note, there were several animals that showed no detectable inhibitory activity against older clades at all (**Fig. 4b**), suggesting that the HA antigen of the currently circulating clade 2.3.4.4b differs significantly from earlier clades, which justified our updating of the vaccine.

**Fig. 4.**
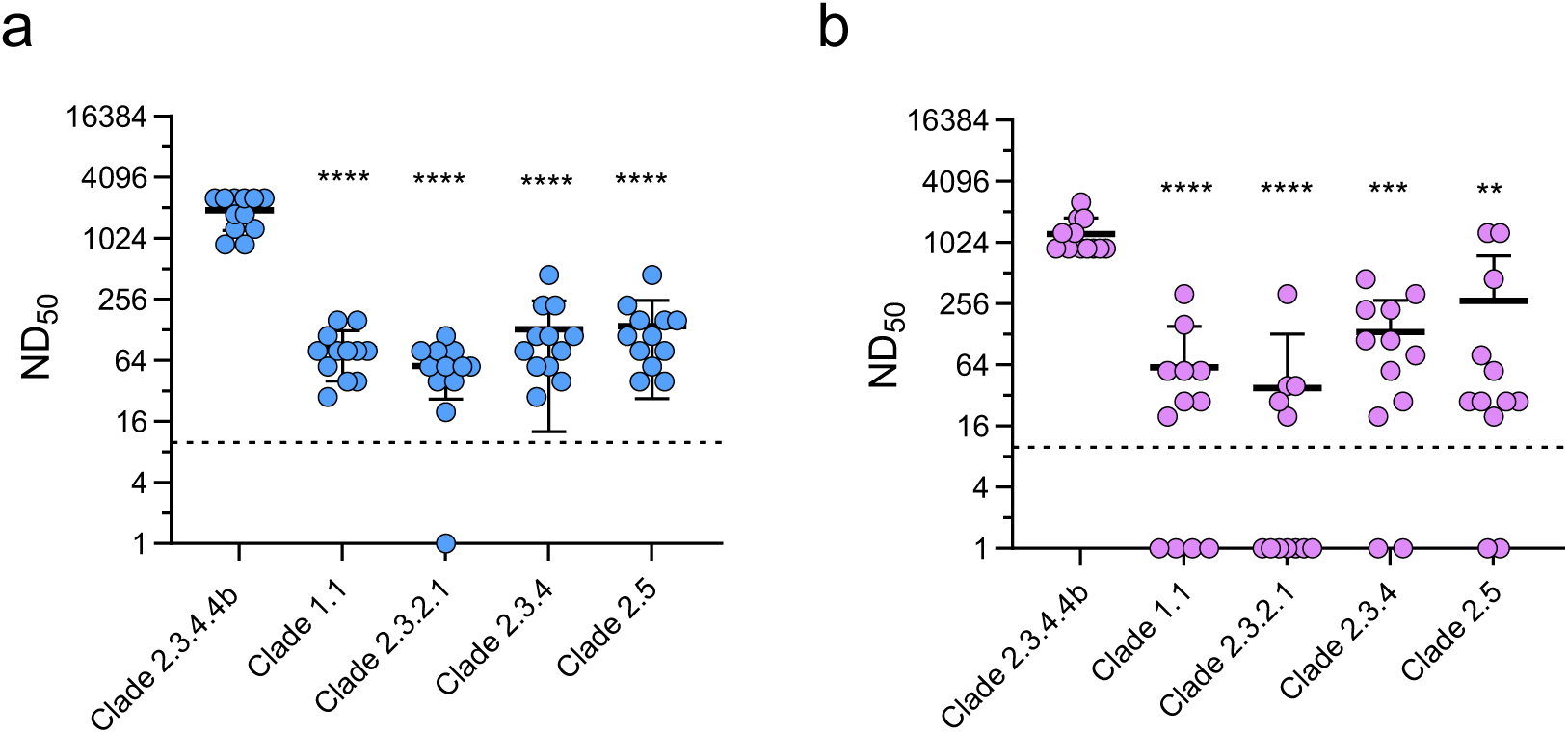
Potency of vaccine-induced neutralizing antibodies against previous H5 clades. (**a, b**) Immune sera were prepared 5 weeks after the second immunization (day 70) from Silky chickens (*Gallus gallus domesticus*, n = 12) (**a),** and African penguins (*Spheniscus demersus*, n = 12) (**b)**, and analyzed for H5N1-neutralizing activity taking advantage of VSVΔG(H5:N1:GFP) surrogate viruses encoding the HA antigens of the indicated H5 clades. Mean values, SD, and lower detection limit are indicated. Significantly different ND_50_ values were computed using the one-way ANOVA with Tukey’s multiple comparison test (**p < 0.01, *** < 0.001, ****P < 0.0001).

## Discussion

In the last decades, numerous countries worldwide have prohibited the general prophylactic vaccination of poultry against H5 and H7 HPAIV. There are several reasons for this non-vaccination policy. First, live-attenuated influenza vaccines (LAIV) based on low-pathogenic H5 or H7 viruses are considered unsafe as they may evolve to HPAIV. Second, inactivated H5 or H7 influenza vaccines that are administered via the parenteral route are regarded as safe, however, it is commonly feared that poultry immunized with inactivated influenza vaccines may appear healthy when infected with HPAIV, but may still shed infectious virus, resulting in unnoticed spread of HPAIV. Third, inactivated influenza vaccines usually do not allow simple serological differentiation between infected and vaccinated animals. However, the DIVA principle is mandatory for efficient control of HPAIV outbreaks, and use of vaccines which do not comply with this principle may lead to trade restrictions ^47^. The current strategy to mitigate HPAIV outbreaks in non-vaccinating countries is based on surveillance, stamping out, and mandated quarantine of poultry and captive wild birds in affected areas ^48^. The non-vaccination policy was sufficiently feasible as long as outbreaks of HPAIV occurred only sporadically and could be controlled by stamping out and quarantine measures. The development of modern vaccines that match the criteria listed above was not a priority up to now. However, in face of the current panzootic by H5N1 HPAIV of clade 2.3.4.4b modern and effective vaccines are missing.

To overcome the hurdles associated with conventional inactivated vaccines, we previously developed an RNA replicon particle vaccine encoding the HA antigen of an H5N1 HPAIV (clade 2.5) isolated in 2004 ^49^. This propagation-defective vector vaccine elicited a strong nAb response in chickens and fully protected the animals from lethal HPAIV infection ^43^. To avoid antigenic mismatch with currently circulating H5N1 HPAIV, we generated in the present study RNA replicon particles encoding the HA of A/Pelican/Bern/1/2022 (H5N1), a clade 2.3.4.4b virus, which had been isolated from a diseased Dalmatian pelican in Bern Animal Park in 2022. The updated VRP vaccine did not replicate in an autonomous way, in line with previous results ^46^, but could be propagated to high infectious titers on helper cells providing the VSV G protein *in trans* ^44^.

Evaluation of this vaccine in SPF chickens showed that a single intramuscular injection of VSVΔG(H5_mb_) replicon particles elicited significant levels of virus-neutralizing serum antibodies. Of note, antibody titers increased significantly after the second administration, indicating that the same vaccine can be used to significantly boost the immune response. The immunized chickens were fully protected against infection with a lethal dose of the autologous H5N1. Moreover, they did not shed H5N1 HPAIV, in line with the previous notion that sterile immunity may be achieved with these VRP vaccines ^43^.

Previous vaccination trials in zoo birds using inactivated, adjuvanted influenza vaccines revealed that various avian species responded differentially to immunization ^40,50–52^. In contrast, the majority of the 24 avian species in the present study responded well to vaccination with adjuvant-free VSVΔG(H5_mb_) replicon particles and produced H5N1-neutralizing antibodies at high levels. One reason for the high efficacy of the VSVΔG(H5_mb_) vaccine in various avian species may be linked to the VSV G protein which was used for *trans*-complementation of the replicon particles. This viral envelope glycoprotein is known to mediate efficient virus entry into a broad range of cell types and tissues of various mammalian species by binding to members of the low-density lipoprotein (LDL) receptor family ^53,54^. Avian homologs of the mammalian LDL receptor family may serve as cellular receptors for VSV however, in Zebra finches LDL receptor homologs were found to be highly divergent and only inefficiently recognized by lentiviruses pseudotyped with the VSV G protein ^55^. LDL receptor diversity may also exist in other avian species, and this might explain why some species enrolled in our study, e.g. the Ferruginous duck *(Aythya nyroca*), the Common goldeneye (*Bucephala clangula*), and the Barnacle goose (*Branta leucopsis*), responded less efficiently to immunization with VSVΔG(H5_mb_).

In contrast to other virus-vectored vaccines for poultry that are based on either Newcastle disease virus ^56,57^, avian herpesviruses ^58–61^, or fowlpox virus ^62^, VSV is not a known avian pathogen. Therefore, birds do not have pre-existing immunity to VSV which could interfere with vaccine efficacy. However, nAb directed to the VSV G protein were detected following immunization with VSVΔG(H5_mb_). In particular, Dalmatian pelicans responded to vaccination with VSVΔG(H5_mb_) by also producing VSV-specific nAbs. This may explain why these animals did not respond well to the second immunization. One year after the second immunization, however, when VSV-specific nAbs titers had dropped, the animals responded to the third immunization. Interestingly, some zoo birds as for example the Greater flamingos showed pre-existing immunity to H5Nx, suggesting that they must have been in contact to subtype H5 AIV prior to vaccination. Despite this pre-existing H5-specific immunity, the flamingos responded very well to vaccination with VSVΔG(H5_mb_), indicating that pre-existing H5-specific nAbs do not interfere with VSVΔG(H5_mb_) infection.

With the exception of birds with previous contact to H5Nx viruses, infected animals were easily discriminated from vaccinated ones using a commercially available ELISA. Thus, the VSVΔG(H5_mb_) vaccine meets two important criteria which are in favor of general vaccination of poultry against HPAIV: (1) The vaccine induces a strong immune response capable of completely blocking shedding of H5N1 HPAIV from vaccinated animals, and (2) it fully complies with the DIVA principle. We also found that vaccination with VSVΔG(H5_mb_) encoding the HA antigen of currently circulating H5N1 HPAIV (clade 2.3.4.4b) elicited serum antibodies that neutralized H5N1 clades isolated about 20 years ago. We therefore hypothesize that the VRP vaccine will also protect birds from antigen-drifted H5N1 which may evolve in the next couple of years.

Another interesting feature of the VSVΔG(H5_mb_) vaccine is the long-lasting immune response that was induced in captive wild birds by this vaccine. One year after the prime/boost vaccination, the majority of animals still showed relatively high virus-neutralizing antibody titers that can be regarded as protective. In Silky chickens, the nAb levels were still high one year after the second immunization, although the animals did not experience any infection by H5Nx viruses which could have boosted the vaccine-induced immune response. Other species such as Greater flamingos and African penguins showed only a moderate drop of nAbs after one year. Thus, vaccination with VSVΔG(H5_mb_) can elicit a long-lasting immune response in various avian species, so that booster immunizations will only be necessary at longer intervals. Furthermore, long-term housing of captive birds indoors may be redundant and thus animal welfare may be improved.

In the present study, we evaluated the VSVΔG(H5_mb_) vaccine in captive wild birds, however, this vaccine might also be used to protect highly endangered avian species *in situ* such as California condors (*Gymnogyps californianus*) and African penguins ^63^. Since one single immunization was sufficient to induce significant nAb levels in most avian species, use of VSVΔG(H5_mb_) may eliminate the need for a second immunization, which is regarded as impractical for free-living wild birds. In line with our previous work ^42,43^, VSV-derived RNA replicon particles also elicit a protective immune response in domestic chickens, recommending this DIVA-compatible vaccine for use in poultry as well. Vaccination of poultry not only would avoid the huge economical losses that outbreaks by H5N1 HPAIV can cause, it would also reduce the risk of zoonotic H5N1 transmission to humans, as exemplified by the vaccination campaign against H7N9 in China ^64^.

In addition to avian species, influenza vaccines based on VSV-derived replicon particles have been proven to work in mammalian species as well ^65–67^. The VSVΔG(H5_mb_) vaccine may be useful for the protection of mammalian species that are affected by the H5N1 HPAIV panzootic, such as dairy cows ^34^, minks ^34,68^, and pet animals such as cats ^27^.

In conclusion, we developed an extraordinary safe and efficacious DIVA vaccine for the protection of chickens and various wild bird species against H5N1 HPAIV. This vaccine may help to control the current H5N1 panzootic and to reduce the risk of zoonotic transmission of H5N1 to humans.

## Materials and Methods

### Cells

Madin-Darby canine kidney (MDCK, type II) cells were kindly provided by Georg Herrler (University of Veterinary Medicine, Hannover, Germany) and grown with Earle’s Minimal Essential Medium (MEM, Life Technologies, Zug, Switzerland) supplemented with 5% fetal bovine serum (FBS, Pan Biotech, Aidenbach, Germany). BHK-21 cells were purchased from the German cell culture collection (DSZM, Braunschweig, Germany) and grown in Glasgow’s Minimum Essential Medium (GMEM, Life Technologies). BHK-G43 cells, a transgenic baby hamster kidney (BHK-21) cell clone producing the VSV G protein in a regulated manner ^44^, were cultured with GMEM supplemented with 5% FBS. Vero E6 cells were kindly provided by Christian Drosten and Marcel Müller (Charite, Berlin, Germany) and maintained in Dulbecco’s minimal essential medium (DMEM; Life Technologies) supplemented with 10% FBS and non-essential amino acids (Life Technologies).

### Viruses

A/Pelican/Bern/1/2022 (H5N1) was isolated from biopsy material prepared from a diseased Dalmatian Pelican in Bern Animal Park in February 2022. Briefly, brain and kidney tissues were homogenized, subject to centrifugation (5 minutes at 14.000 x g, 4°C), and the clear supernatant sterile-filtrated (Minisart NML cellulose acetate standard syringe filter with 0.2 µm pore size). The material passing the filter was inoculated with confluent MDCK monolayers for 2 days at 37°C. The supernatant was collected, cell debris removed by centrifugation, and the clear supernatant supplemented with 10% FBS, and stored in aliquots at -70°C. Infectious virus titers were determined on MDCK cells by limiting dilution. The infected cells were washed once with PBS and fixed with 10% formalin containing 1% (w/v) of crystal violet. The tissue infectious dose 50% (TCID_50_) was calculated according to the Spearman-Kärber formula ^69^.

VSV*, a propagation-competent VSV encoding the green fluorescent protein (GFP) reporter protein, and the propagation-defective VSV*ΔG which encodes the GFP reporter protein but lacks the G protein gene have been described previously ^70^. Both viruses were titrated on BHK-21 or Vero cells as described ^70^.

The propagation-competent chimeric viruses VSVΔG(HA_Y_:NA_Y_:GFP) encoding the HA and NA proteins of A/chicken/Yamaguchi/7/2004 (H5N1), clade 2.5, VSVΔG(HA_V_:NA_Y_:GFP) encoding the HA protein of A/Muscovy duck/Vietnam/OIE-559/2011 (H5N1), clade 1.1, and VSVΔG(HA_HK_:NA_Y_:GFP) encoding the HA protein of A/peregrine falcon/Hong Kong/810/2009 (H5N1), clade 2.3.4, and VSVΔG(HA_Ho_:NA_Y_:GFP) encoding the HA protein of A/whooper swan/Hokkaido/4/2011 (H5N1), clade 2.3.2.1, have been described in a previous report ^46^. These chimeric reporter viruses were propagated and titrated on MDCK cells.

### Plasmids

Viral RNA was isolated with TRIzol reagent (Life Technologies, cat. no. 15596026) from 200 μl of A/Pelican/Bern/1/2022 (H5N1) stock (passage 2 on MDCK cells) according to the manufacturer’s instructions. The RNA was reverse-transcribed by the SuperScript III Reverse Transcriptase (Life Technologies, cat. no. 18080093) using the Uni12 primer that is complementary to the conserved 12 nucleotides of the 3’-end of the viral RNA ^71^. Segment 4 and segment 6 cDNAs were amplified with Phusion™ High-Fidelity DNA Polymerase (Life Technologies, cat. no F530L) using segment-specific sense and antisense primers ^72^. The PCR products were gel-purified, cloned into the pJET1.2 plasmid (Life Technologies, cat. no. K1231), and sequenced according to Sanger. The cDNA sequences of segment 4 and segment 6 have been deposited at the GISAID databank (accession nos. EPI3526757 and EPI3526758, respectively).

The open reading frame of A/Pelican/Bern/1/2022 (H5N1) HA was amplified with Phusion high fidelity DNA polymerase and cloned into the *Mlu*I and *Nhe*I sites of the pVSV*ΔG(HA) plasmid ^42^, resulting in plasmid pVSVΔG(H5_pb_). The HA polybasic (pb) proteolytic cleavage site REKRRKR was changed into a monobasic (mb) sequence motif (RETR) by taking advantage of an overlapping PCR method ^73^. The modified H5 cDNA was subsequently amplified and cloned into the pVSV*ΔG(HA) plasmid resulting in pVSVΔG(H5_mb_). To generate a propagation-competent chimeric VSV reporter virus, the HA, and NA genes of A/Pelican/Bern/1/2022 (H5N1) were inserted into the *Mlu*I and *Xho*I sites of pVSVΔG(HA_R_:NA_R_:GFP) ^74^ by In-Fusion cloning (In-Fusion Snap Assembly Master Mix, Takara, Saint-Germain-en-Laye, France, cat. no. 638948), resulting in the plasmid pVSVΔG(HA_P_:NA_P_:GFP).

### Generation of recombinant RNA replicon particles

Virus replicon particles (VRPs) have been produced according to a published procedure ^43^. Briefly, BHK-G43 cells were inoculated for 1 hour with modified vaccinia virus Ankara encoding the T7 phage RNA polymerase (MVA-T7) ^75^, using a multiplicity of infection (m.o.i.) of 3 focus-forming units (f.f.u.) per cell. Subsequently, the cell culture medium was replaced with GMEM containing 5% FBS and 10^-9^ M mifepristone (Merck KGaA, Darmstadt, Germany), and the cells transfected with the Lipofectamine 2000 transfection reagent (Life Technologies) and a mixture of four plasmids: pVSVΔG(H5_mb_:N1:GFP (or pVSVΔG(H5_pb_:NA:GFP) and plasmids encoding the VSV N, P, and L genes, respectively, all under the control of the T7 promotor (Kerafast, Boston, USA; cat. no. EH1012). One day post transfection, the cells were dissociated with trypsin/EDTA (Life Technologies) and co-cultured for 24 hours with an equal number of non-transfected BHK-G43 cells in the presence of 10^-9^ M mifepristone. The cell culture supernatant was collected and cell debris removed by centrifugation (1200 x g, 10 minutes, 4°C). The clarified cell culture supernatant was passed through a 0.2 µm pore size filter for depletion of MVA-T7. The replicon particles VSVΔG(H5_mb_) and VSVΔG(H5_pb_) were propagated on mifepristone-treated BHK-G43 cells and stored in aliquots at -70 °C in the presence of 10% FBS.

### Titration of RNA replicon particles

Infectious virus titers were determined on BHK-21 cells grown in 96-well microtiter plates. The cells were inoculated in duplicate with 40 µl per well of serial 10-fold virus dilutions for 1 hour at 37 °C. Thereafter, 60 µl of GMEM was added to each well, and the cells incubated for 24 hours at 37 °C. The cells were fixed for 30 minutes with 3.7% formalin in PBS, permeabilized with 0.25% (v/v) of Triton X-100, and incubated for 60 minutes with a monoclonal antibody directed to the VSV matrix protein (mAb 23H12, diluted 1:25 with PBS, KeraFast, Boston, MA, cat. no. EB0011) and subsequently for 60 minutes with goat anti-mouse IgG conjugated with Alexa Fluor-488 (diluted 1:500 in PBS; Life Technologies, cat. no. A28175). Infected cells were detected by fluorescence microscopy (Observer.Z1 microscope, Zeiss, Feldbach, Switzerland), and infectious virus titers were calculated and expressed as focus-forming units (f.f.u.)/ml.

### Preparation of the RNA replicon vaccine

For a typical vaccine batch preparation, BHK-G43 cells were seeded into five T150 flasks and maintained in 40 ml/flask of GMEM medium with 5% FBS. When confluency was reached, the medium was aspirated, and cell maintained at 37°C for 6 hours with 40 ml/flask of GMEM containing 10^-9^ M of mifepristone. VSVΔG(H5_mb_) was added (m.o.i of 0.1 f.f.u./cell), and cells further incubated at 37°C for 20 hours. The cell culture supernatant was transferred to 50-mL Falcon tubes and cell debris removed by centrifugation (1000 x g, 15 min, 4°C). The replicon particles in the cleared supernatant were pelleted by ultracentrifugation (105’000 x g, 60 min, 4°C), and resuspended in 20 mL of PBS. The VSVΔG(H5_mb_) replicon vaccine was stored in 5-mL aliquots at -70°C.

### Western blot analysis

Confluent monolayers of BHK-G43 cells grown in T75 flasks were treated for 6 hours with mifepristone (10^−9^ M) and subsequently infected with either VSVΔG(H5_pb_) or VSVΔG(H5_mb_) using an m.o.i. of 0.1 f.f.u./cell. At 24 hours p.i., the cell culture supernatant of the infected cells was collected and cell debris removed by centrifugation (1200 x g, 15 min, 4°C). Subsequently, virus particles were pelleted from the clarified cell culture supernatant by ultracentrifugation and solubilized by adding preheated (95°C) sodium dodecyl sulfate (SDS) sample buffer containing 0.1M of dithiothreitol (DTT) to the pellets. The solubilized proteins were separated by SDS polyacrylamide gel electrophoresis (PAGE) using two 4–12% gradient gels (SurePAGE™; Genscript, Leiden, The Netherlands). The separated proteins were visualized by incubating the first gel overnight with colloidal Coomassie (GelCode Blue Stain Reagent, Life Technologies, cat. no. 24590). The separated proteins of the second gel were transferred to a nitrocellulose membrane by semi-dry blotting. The nitrocellulose membrane was blocked overnight at 4 °C with Odyssey Blocking Reagent (Li-COR Biosciences, Lincoln, NE) and subsequently incubated with a polyclonal chicken immune serum which was directed against the low-pathogenic A/duck/Hokkaido/Vac-1/2004 (H5N1) ^43^. The membrane was washed four times with PBS containing 0.1% Tween-20 and incubated with the secondary antibodies IRDye 800CW donkey anti-chicken IgY (LI-COR Biosciences, Cat. no. 926-32218) diluted 1:5000 in PBS. Following several washing steps with PBS/0.1% Tween 20, the blots were scanned with the Odyssey Infrared Imaging system (LI-COR Biosciences, Bad Homburg, Germany).

### Indirect immunofluorescence analysis

MDCK cells grown on 12-mm glass cover slips were inoculated for 90 minutes with either VSVΔG(H5_pb_), VSVΔG(H5_mb_) or VSVΔG(H5_P_:N1_P_:GFP) using an m.o.i. of 0.02 f.f.u./cell. The cells were washed once with MEM, further incubated for 20 hours at 37°C, and fixed for 30 minutes with PBS containing 3.7% of formalin. Cells that had been infected with VSVΔG(H5_P_:N1_P_:GFP) were directly visualized by fluorescence microscopy taking advantage of the GFP reporter protein. Cells that had been infected with either VSVΔG(H5_pb_) or VSVΔG(H5_mb_) were permeabilized for 5 minutes with 0.25% (vol/vol) of Triton X-100 in PBS, incubated for 1 hour with a monoclonal antibody directed to the influenza nucleoprotein (clone H16-L10-4R5; ATCC, HB-65) diluted 1:50 in PBS, and subsequently incubated for 1 hour with goat anti-mouse IgG conjugated to Alexa Fluor-488 (1:500; Life Technologies, cat. no. A11001). Finally, the cells were embedded in Mowiol 4–88 medium (Merck KGaA) and analyzed by fluorescence microscopy.

### Animal experiments

This study was conducted in the BSL3-Ag containment facilities of the Institute of Virology and Immunology IVI (Mittelhäusern, Switzerland), in compliance with Swiss animal welfare regulations (TSchG SR 455; TSchV SR 455.1; TVV SR 455.163). The committee on animal experiments of the canton of Bern, Switzerland, reviewed the experiments, and the cantonal veterinary authority (Amt für Landwirtschaft und Natur LANAT, Veterinärdienst VeD, Bern, Switzerland) approved the study under the authorization numbers BE24/2023 and BE26/2023. Specific pathogen-free (SPF) White Leghorn chickens (5 to 6 weeks old, mixed sex) were obtained from the IVI breeding flock. Animals A, B, and C (group size n = 8) were immunized intramuscularly (i.m.) by injecting 250 μl of either VSV*ΔG (group A) or VSVΔG(H5_mb_) (groups B and C) into the left and the right breast muscle (500 μl in total, corresponding to 10^8^ f.f.u./animal). Animals of group D were immunized with VSVΔG(H5_mb_) by dropping 100 μl of VRP suspension into each eye (corresponding to 10^8^ f.f.u./animal). The animals were kept for 4 weeks employing deep litter management and ‘‘ad libitum’’ access to feed and water. At day 28, chickens of groups A and B were boosted (i.m.) with 10^8^ f.f.u. of VSV*ΔG and VSVΔG(H5_mb_), respectively, while animals of groups C and D were immunized via eye drop using 10^8^ f.f.u./animal of VSVΔG(H5_mb_). At day 58, the chickens were infected via the nasal route with 50 μl of A/Pelican/Bern/1/2022 (H5N1) containing 10^6^ TCID_50_. Following infection, the animals were surveyed daily for clinical signs of disease. A clinical scoring system was used to define human endpoint criteria. Swab samples were collected at days 1, 2, 3, 5, and 7 p.i. and tested for viral RNA by RT-qPCR (see below). At day 68, all surviving animals were euthanized. Blood was also collected at day 1 (pre-immune), day 28 (prime), day 56 (boost), and day 68 (post mortem). Serum was prepared by centrifugation of coagulated blood and stored at -20°C before analysis by ELISA or virus neutralization test (see below).

For vaccination of captive wild birds, 348 birds belonging to 24 different species were selected in the zoo of Basel and Bern Animal Park. In particular birds living in outdoor enclosures with potential exposure to avian influenza were included in the study. Prior to immunization, all birds were subject to clinical examination including assessment of general health, body condition and body weight. Only birds deemed clinically healthy were included in the study. Blood samples were collected at days 1, 35, 70, 365, and 400 using either the *Vena jugularis*, *Vena ulnaris* or the *Vena metatarsalis medialis* as sampling sites. Approximately 1.0 mL of blood was collected per bird using 22G to 27G disposable hypodermic needles (100 Sterican®, B. Braun Melsungen AG, Germany), depending on the size of the bird. A total of 317 birds were vaccinated three times (at days 1, 35, and 365) using 10^8^ f.f.u./mL of VSVΔG(H5_mb_). The vaccine was usually administered by injection into the *M. pectoralis*. However, in case of the pelicans (*Pelecanus crispus* and *Pelecanus onocratus*), which have subcutaneous air sacs with large extensions, *M. gastrocnemius* was used for injection. Birds with body weights of less than 1.5 kg received 0.25 mL of the vaccine (corresponding to 5x 10^7^ f.f.u./animal.), while birds with more than 1.5 kg of body weight received 0.5 mL of the VRP suspension. Following immunization, zoo birds were closely monitored for seven days by the attending veterinarians and animal caretakers in order to detect potential adverse reactions or changes in animal behavior in response to the vaccination.

### RT-qPCR

To determine virus loads in oropharyngeal and cloacal swab samples, swab tips were placed into 0.5 mL of RA1 lysis buffer (Macherey-Nagel, Düren, Germany; cat. no. 740961) containing 1% (v/v) of β-mercaptoethanol. After a short centrifugation step, RNA was extracted from 200 μl of lysate using the NucleoMag Vet kit (Macherey Nagel, cat. no. 744200) according to the manufacturer’s protocol. Reverse transcription from RNA to cDNA and real-time quantitative PCR (qPCR) were performed on the QuantStudio 5 real-time PCR system (Thermo Fisher Scientific) using the AgPath-ID One-Step RT-PCR kit (Life Technologies, cat. no. AM1005) and segment 7-specific oligonucleotide primers and probe ^76,77^. Data were acquired and analyzed using the Design and Analysis Software v1.5.2 (Thermo Fisher Scientific).

### Enzyme-linked Immunosorbent Assay (ELISA)

For detection of serum antibodies directed to the influenza nucleoprotein (NP) and the H5 hemagglutinin, the ID Screen® Influenza A Antibody Competition Multi-species ELISA (ID-Vet, Montpellier, France, cat. no. FLUACA) and the ID Screen® Influenza H5 Antibody Competition (ID-Vet, cat. no.: FLUACH5-5P) were used, respectively. ELISA were performed according to the manufacturer’s instructions.

### Virus neutralization tests

Serum samples were heated for 30 minutes at 56°C to inactivate factors of the complement system. Twofold serial dilutions of heat-inactivated immune sera were prepared in duplicates or quadruplicates in 96-well cell culture plates using MEM cell culture medium (50 µl/well). To each well, 50 µl of cell culture medium containing 100 f.f.u. of either VSVΔG(H5_P_:N1_P_:GFP) or VSV-GFP were added and incubated for 30 min at 37°C. Subsequently, the antibody/virus mix was transferred to confluent MDCK or Vero cell monolayers in 96-well cell culture plates and incubated at 37°C for 24 hours. Infected cells were detected by fluorescence microscopy taking advantage of the virus-encoded GFP reporter. Neutralization doses 50% (ND_50_) values were calculated according to the Spearman and Kärber method ^69^.

Neutralization tests with authentic A/Pelican/Bern/1/2022 (H5N1) were performed analogously. However, the assay was performed in quadruplicates using 100 TCID_50_ of virus per well. The antibody/virus mixtures were incubated with MDCK cells for 48 hours. Thereafter, the cells were washed once with 200 μl of PBS/well and fixed for 60 minutes at 21°C with 10% formalin containing 1% crystal violet. Finally, the microtiter plates were rinsed with tape water and the ND_50_ titer calculated as above.

### Statistical analysis

Statistical analysis was performed using GraphPad Prism 10, version 10.1.2 (GraphPad Software, Boston, Massachusetts, USA). Unless noted otherwise, the results are expressed as mean ± SD. Specific statistical tests such as the one-way or two-way ANOVA test were used to assess significant differences in serum antibody responses in vaccinated animals as indicated in the figure legends. *P* values < 0.05 were considered as significant.

### Biosafety

Work with HPAIV A/Pelican/Bern/1/2022 (H5N1) has been approved by the Swiss Federal Office of Public Health (license no. A230074-01) and was performed at IVI laboratories and animal facilities complying to biosafety level 3. Release of the genetically modified virus replicon vaccine was approved by the Swiss Federal Office for the Environment (FOEN) (reference no. BAFU-217.2344635/2) with consent of the Federal Office of Public Health (FOPH), the Federal Food Safety and Veterinary Office (FSVO), and the Federal Office for Agriculture (FOAG).

## Acknowledgements

This work received financial support from the Foundation Tierspital in Basel. The funders had no role in study design, data collection and analysis, decision to publish or preparation of the manuscript. We thank the animal caretakers Katarzyna Sliz and Daniel Brechbühl at IVI for their assistance in experimental infection of vaccinated chickens. We are grateful to Hansueli Blatter and his team of animal caretakers at Bern Animal Park and all the animal caretakers at the zoo of Basel for their excellent assistance in animal care and handling. We thank Sandra Renzullo, Yelena Ruedin, and Markus Mader from the diagnostic team at IVI for practical laboratory support. We gratefully acknowledge the scientific support of Claudia Bachhofen (IVI), Barbara Wieland (IVI), and Friederike von Houwald (Bern Animal Park) and veterinary assistance by Seraina Meister (Zoo Basel) and Michael Rüttener (Zoo Basel).

