## Supplementary Fig. 1 for "Immunization with a novel RNA replicon vaccine confers long-lasting protection against H5N1 avian influenza virus in 24 bird species"

```
                             1                                           ------------------
HA1-2.3.4.4b                 DQICIGYHANNSTEQVDTIMEKNVTVTHAQDILEKTHNGKLCDLNGVKPLILKDCSVAGW
HA1-2.3.4                    DQICIGYHANNSTEQVDTIMEKNVTVTHAQDILEKTHNGKLCDLDGVKPLILRDCSVAGW
HA1-2.3.2.1                  DHICIGYHANNSTEQVDTIMEKNVTVTHAQDILEKTHNGKLCDLNGVKPLILKDCSVAGW
HA1-2.5                      DQICIGYHANNSTEQVDTIMEKNVTVTHAQDILEKTHNGKLCDLDGVKPLILRDCSVAGW
HA1-1.1                      DQICIGYHANNSTEQVDTIMEKNVTVTHAQDILEKTHNGKLCDLDGIRPLILRDCSVAGW

                            61----------------------------------------------------------
HA1-2.3.4.4b b               LLGNPMCDEFIRVPEWSYIVERANPANDLCYPGSLNDYEELKHLLSRINHFEKILIIPKS
HA1-2.3.4                    LLGNPMCDKFNDVPEWSYIVEKTNPANDLCYPGNLNDYEELKHLLSRINHFEKIQIFPKS
HA1-2.3.2.1                  LLGNPLCDEFINVPEWSYIVEKANPANDLCYPGNFNDYEELKHLLSRINHFEKIQIIPKD
HA1-2.5                      LLGNPMCDEFINVPEWSYIVEKANPSNDLCYPGNFNDYEELKHLLSRINHFEKIQIIPKS
HA1-1.1                      LLGNPMCDEFINVPEWSYIVEKANPVNDLCYPGVFNDYEELKHLLSRINHFEKIQIIPKS

                            121---------------------------------------------------------
HA1-2.3.4.4b                 SWPNHETSLGVSAACPYQGAPSFFRNVVWLIKKNDAYPTIKISYNNTNREDLLILWGIHH
HA1-2.3.4                    SWPDHEASLGVSAACPYQGMPSFFRNVVWLIKKNNTYPTIKSSYNNTNKEDLLILWGIHH
HA1-2.3.2.1                  SWSDHEASLGVSAACSYQGNSSFFRNVVWLIKKDNAYPTIKKGYNNTNQEDLLVLWGIHH
HA1-2.5                      SWSDHEASSGVSSACPYQGRSSFFRNVVWLIKKNSAYPTIKRSYNNTNQEDLLVLWGIHH
HA1-1.1                      SWPSHEASLGVSAACPYQGNSSFFRNVVWLIKKNSTYPTIKRSYNNTGQEDLLVMWGIHH
                                   130-loop
                            181---------------------------------------------------------
HA1-2.3.4.4b                 SNNAEEQTNLYKNPTTYISVGTSTLNQRLVPKIATRSQVNGQRGRMDFFWTILKPDDAIH
HA1-2.3.4                    SNNEAEQTDLYQNSNTYISVGTSTLNQRLVPKIATRSKVNGQSGRMDFFWTILKPNDAIN
HA1-2.3.2.1                  PNDEAEQTRLYQNPTTYISIGTSTLNQRLVPKIATRSKINGQRGRIDFFWTILKPNDAIH
HA1-2.5                      PNDAAEQTRLYQNPTTYISVGTSTLNQRLVPKIATRSKINGQSGRMEFFWTILKPNDAIS
HA1-1.1                      PNDAAEQTKLYQNPTTYISVGTSTLNQRLTPRIATRSKVNGQSGRMEFFWTILKPNDAIN
                                  190-helix                          220-loop
                            241------------------------------------
HA1-2.3.4.4b                 FESNGNFIAPEYAYKIVKKGDSTIMKSGVEYGHCNTKCQTPVGAINSSMPFHNIHPLTIG
HA1-2.3.4                    FESNGNFIAPEYAYKIVKKGDSAIMKSEVGYGNCNTKCQTPIGAINSSMPFHNIHPLTIG
HA1-2.3.2.1                  FESNGNFIAPEYAYKIVKKGDSTIMKSEVEYGNCNTRCQTPIGAINSSMPFHNIHPLTIG
HA1-2.5                      FESNGNFIAPEYAYKIVKKGDSAIMKSELEYGNCNTKCQTPMGAINSSMPFHNIHPLTIG
HA1-1.1                      FESNGNFIAPEYAYKIVKKGDSTIMKSELEYGNCNTKCQTPMGAINSSMPFHNIHPLTIG

                            301
HA1-2.3.4.4b                 ECPKYVKSNKLVLATGLRNSPL
HA1-2.3.4                    ECPKYVKSNKLVLATGLRNSPL
HA1-2.3.2.1                  ECPKYVKSNKLVLATGLRNSPQ
HA1-2.5                      ECPKYVKSSRLVLATGLRNSPQ
HA1-1.1                      ECPKYVKSNRLVLATGLRNSPQ
```

**Supplementary Fig. 1. Comparison of the HA$_1$ subunit amino acid sequences of different H5 clades.** The HA$_1$ primary amino acid sequences of A/peregrine falcon/Hong Kong/810/2009 (H5N1), clade 2.3.4 (GenBank acc. no. BAI39636), A/whooper swan/Hokkaido/4/2001 (H5N1), clade 2.3.2.1 (GenBank acc no. BAJ71684), A/chicken/Yamaguchi/7/2004 (H5N1), clade 2.5, (GenBank acc. no. AB166864), A/Muscovy duck/Vietnam/OIE-559/2011 (H5N1), clade 1.1 (GenBank acc. no. BAK39622) were compared with the HA1 primary sequence of A/Pelican/Bern/1/2022 (H5N1), clade 2.3.4.4b (GISAID acc. no. EPI3526757) using CLUSTAL W. The globular head region is indicated by a dashed line on top of the sequences. The 130-loop, 190-helix and 220-loop that are part of the receptor-binding pocket are indicated by a greyish background. Amino acid positions differing from the clade 2.3.4.4b reference sequence are highlighted by yellow background. Amino acid residues that are unique for the clade 2.3.4.4b HA$_1$ primary sequence are marked by bold red letters.
