## Supplementary Fig. 2 for "Immunization with a novel RNA replicon vaccine confers long-lasting protection against H5N1 avian influenza virus in 24 bird species"

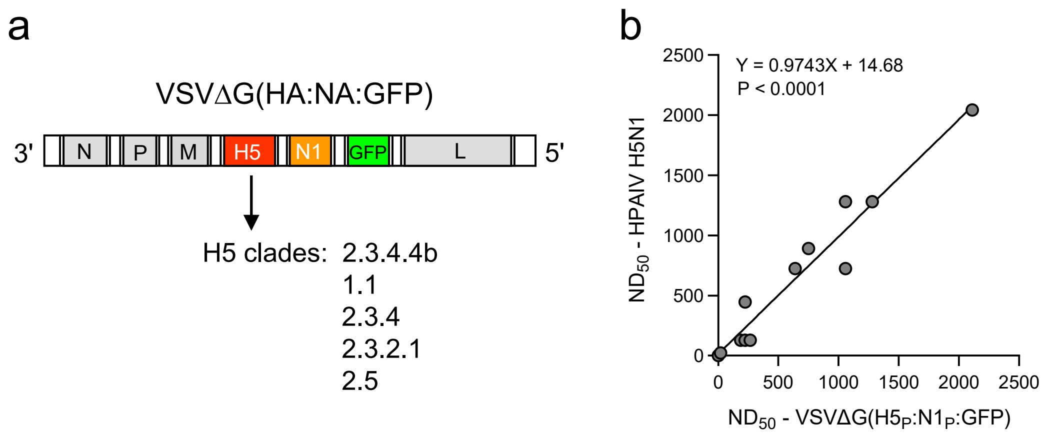


**Supplementary Fig. 2. Comparison of neutralization tests using authentic H5N1 HPAIV and a chimeric VSV reporter virus**. (**a**) Genome map of the propagation-competent VSVΔG(H5:N1:GFP) vector encoding the authentic HA and NA envelope glycoproteins of HPAIV (H5N1) and a GFP reporter. Surrogate viruses encoding the HA antigen derived from either of the indicated H5 clades were used in this study. (**b**) Linear regression analysis of the two virus neutralization tests. Twelve pre-immune and immune sera prepared from the blood of VSVΔG($H5_{mb}$)-immunized chickens were analyzed by virus neutralization tests using either authentic A/Pelican/Bern/1/2022 (H5N1) or VSVΔG($H5_P$:$N1_P$:GFP) encoding the HA and NA genes of this virus isolate. The $ND_{50}$ titers obtained with either of the two assays were analyzed by linear regression.
