## Supplementary Fig. 3 for "Immunization with a novel RNA replicon vaccine confers long-lasting protection against H5N1 avian influenza virus in 24 bird species"

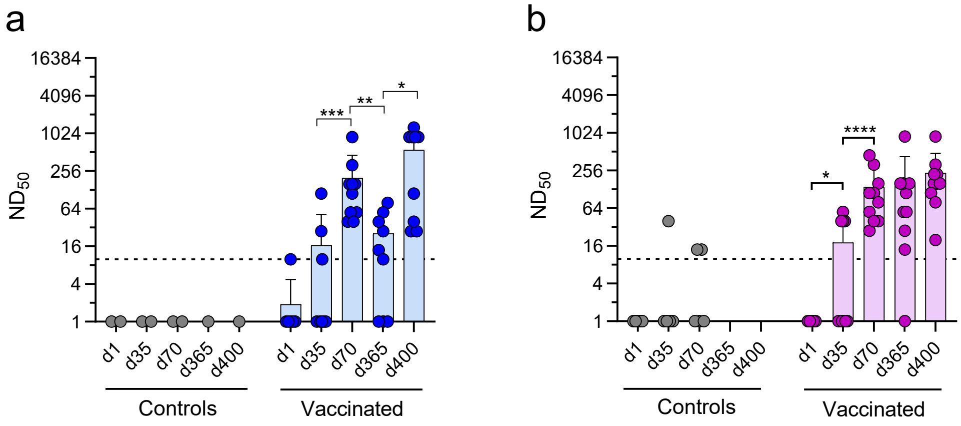


**Supplementary Fig. 3. Detection of VSV-neutralizing antibodies in serum of VSVΔG($H5_{mb}$)-immunized animals. (a, b)** Dalmatian pelicans (*Pelecanus crispus*, n = 10) in the animal park of Bern (a) and Silky chickens (*Gallus gallus domesticus*, n = 10) in the zoo of Basel were immunized intramuscularly with VSVΔG($H5_{mb}$) particles produced on VSV G protein-expressing cells. The animals were boosted with the same vaccine at days 35 and 365. Two Dalmatian pelicans and five Silky chickens were not vaccinated and served as controls. After blood was withdrawn from the animals at the indicated days, sera were prepared and tested for the presence of VSV-specific nAbs using the propagation-competent VSV* reporter virus. Mean $ND_{50}$ values and SD are shown. The dashed line represents the lower detection limit ($ND_{50}$ = 10). The two-way ANOVA test was used to assess significantly different $ND_{50}$ titers between the different time points of blood collection (*$p < 0.05$, **$p < 0.01$, ***$p < 0.001$, ****$p < 0.0001$).
